# Fear not: avoidance behavior missing in species sympatric with the critically endangered West African lion

**DOI:** 10.1101/2020.06.09.142604

**Authors:** Nyeema C. Harris, Kirby L. Mills

## Abstract

Predators are major regulators in communities through trophic and non-trophic pathways. However, as human pressures continue to threaten apex predators, including Africa’s iconic lion, predators’ functional role in their ecosystems may be compromised. Where lions are critically endangered, we found no evidence of avoidance behavior in either competitor or prey species from a camera survey in the largest protected area complex in West Africa. Our findings raise concerns that lions have already become functionally extinct in portions of their West African range, highlighting the urgency of restorative efforts and environmental investments to reverse current declining population trends and the loss of regulatory roles.

## 1. Introduction

African lions face a questionable future. As iconic symbols of their savanna ecosystems, few geographic areas remain of sufficient size and prey abundance to support healthy populations [1]. Where lions do reside, they exist in a persistently threatened state, as almost all populations are declining [2]. In West Africa, lions remain in only 1.1% of their historical range [3]. Alarming rates of environmental change throughout Africa coupled with ongoing human persecution hinder their recovery and expansion [4]. Furthermore, projections of doubling human populations, changing climates, and demands for livestock products on the continent will exacerbate the fragility of extant African lion populations [5-7]. Given shrinking range distributions and declining population densities, the community-regulating pressures normally exerted by lions on sympatric species can be expected to wane. Trophic cascade theory predicts that the regional and global extinction of apex predators can (and indeed often does) have consequences for whole ecosystems [8-10]. For example, savannas without apex predators can harbor sicker animals [11], less diverse plant assemblages [12], and reduced efficiency in nutrient cycling [13]. However, since these pressures are both density dependent and nonlinear, loss of apex predators’ function can precede extirpation.

Non-trophic, trait-mediated effects of apex predators are pervasive and frequently powerful in structuring communities. Animals adapt their behavior to the landscape of risks [14, 15]. For example, ungulates alter habitat selection, foraging patterns, space use, and diel activity to evade their predators [16-19]. Previous studies highlight how heterogeneity of space use along a risk axis influences vegetation patterns (e.g., distribution of thorny *Acacia* [20]) and ecosystem processes (e.g., carbon storage and nutrient cycling [21]) through modulation of the distribution of grazing and browsing pressures by ungulates. Carnivores also modify their behavior to evade dominant competitors to reduce competition and direct mortality [22, 23]. For example, cheetahs (*Acinonyx jubatus*) have been observed to abandon prey and reduce prey handing times in the presence of lions [24]. Despite much research on African ecosystems and apex predation regulation, our understanding of these non-trophic effects in regions where lion populations have dwindled to near extinction remains limited. The loss of lions as effective regulators of carnivore and herbivore communities stands to transform entire West African savanna ecosystems.

In the present study, we assess dampening of fear effects via avoidance behavior exerted by the critically endangered West African lion (figure 1). As giving-up density experiments and radio-telemetry data are scant, our work provides a precursory understanding of changing non-trophic dynamics where lions are now rare. Specifically, we quantify differential patterns of space use in hyenas (*Crocuta crocuta*) and six common prey species corresponding to areas of low and high intensity of use by lions via occupancy models. If West African lions continue to function as system regulators, exerting top-down and intraguild pressures to change behavior within the community, we expect lion occupancy to deter space use by both competitor and prey species. As a result, hyenas and ungulates should exhibit reduced occupancy in areas heavily frequented by lions.

**Figure 1.**
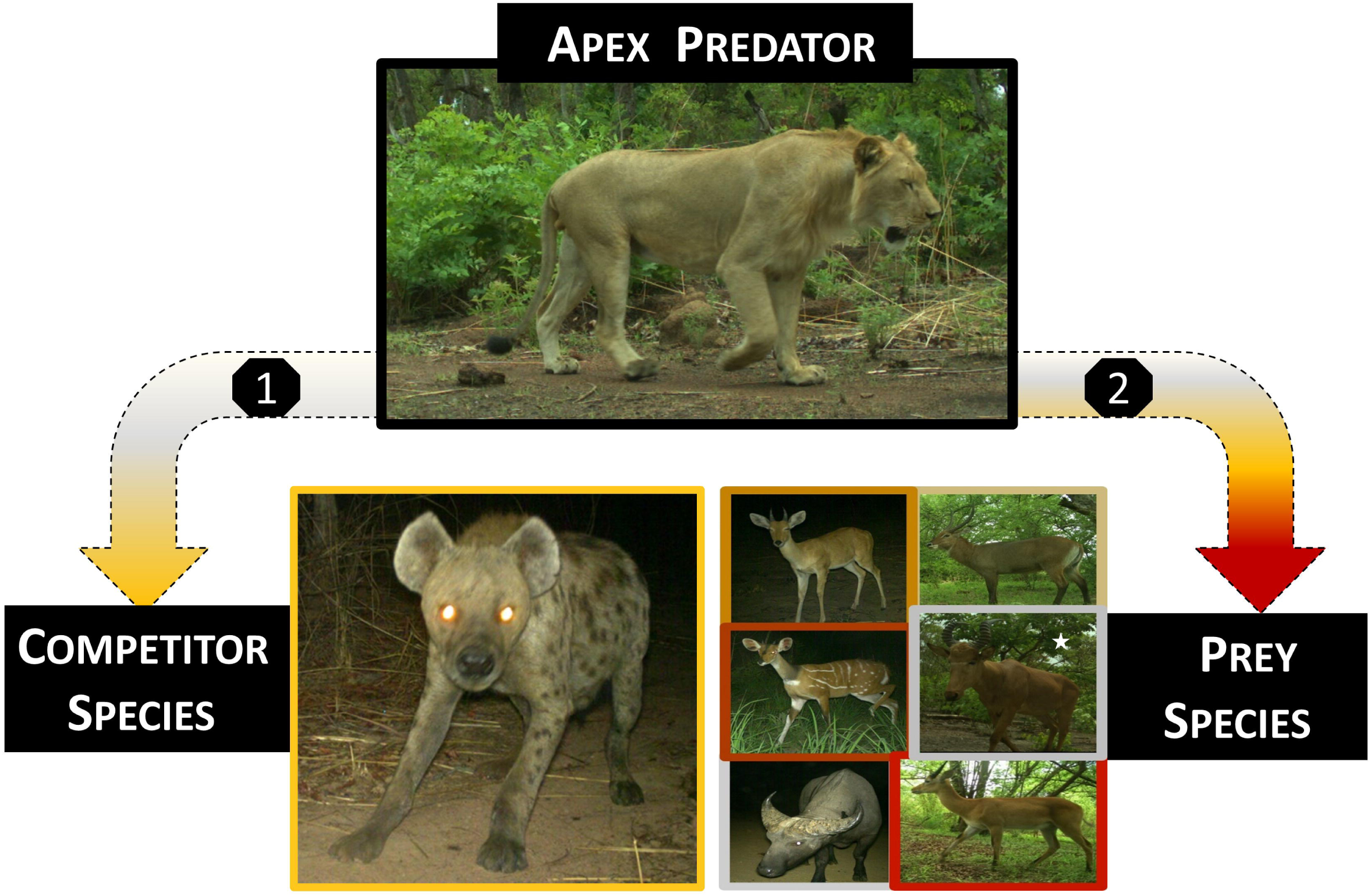
Non-trophic pathways from apex predator on sympatric species, example from African ecosystem. 1) African lions on hyenas as a dominant competitor. 2) African lions on ungulate as prey resources: reedbuck (*Redunca redunca*), waterbuck (*Kobus ellipsiprymnus*), bushbuck (*Tragelaphus sylvaticus*), hartebeest (*Alcelaphus buselaphus*), buffalo (*Syncerus caffer*), kob (*Kobus kob)* - by row. Gradient in arrows reflects a greater effect of lions on prey than the competitor species. Color of border on images indicates hypothesized differential effects of intensity from apex predator on competitor and prey species (darker colors = higher intensity).

## 2. Material and Methods

### (a) Study area and field methods

We implemented a large scale, non-invasive survey, installing 238 camera stations across 21,480 trap night in the W-Arly-Pendjari (WAP) transboundary protected area complex from 2016-2018 (figure 2a). WAP comprises 26,515 km2 of mainly national parks and hunting concessions in portions of Burkina Faso, Niger, and Benin and was inscribed as a UNESCO World Heritage site in 1996. We divided the study area into 10 x 10 km grid cells and systematically deployed cameras within 2 km of the centroid in each grid cells. Unbaited cameras were placed ∼0.5 m off the ground, affixed to trees > 0.5m diameter with microsite selection based on animal sign such as tracks, scat, game trails, and vegetation disturbance. We conducted our camera survey from February to June of 2016-2018 in the national parks and hunting concessions of Burkina Faso and Niger, comprising 49% of the WAP transboundary complex of West Africa. See Harris *et al*. [25] and Mills *et al*. [26] for more details on survey effort and image processing.

**Figure 2.**
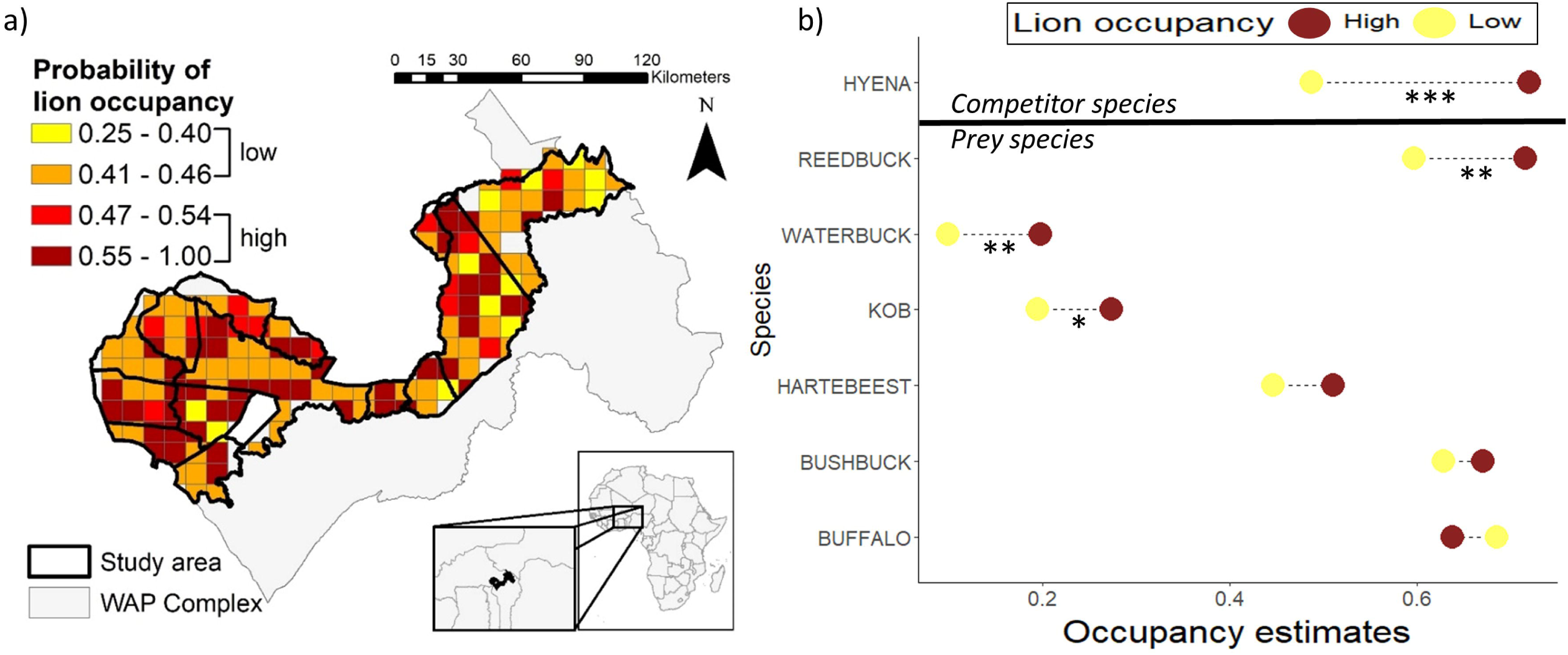
a) Lion occupancy estimates per 10 x 10 km grid cell within study area located in the W-A-P transboundary complex of West Africa. Estimates obtained via camera survey from Mills *et al*. 2020. b) Change in mean occupancy for competitor and prey species corresponding to zones of high and low lion occupancy. Low (n=88 grids) and high (n=116 grids) zones defined as lion occupancy below and above 0.464, the median lion occupancy in 10 x10 km grids, respectively. Mann-Whitney test for significance: * p < 0.05, ** p < 0.01, *** p < 0.001.

### (b) Occupancy modeling

From images identified and confirmed as focal study species, we built single-species, single season occupancy models to evaluate space use of hyena and six prey species in relation to lion occupancy. Detection/non-detection data for each grid cell were separated into 2-week survey periods to model cell-specific probabilities of detection (*p*) and occupancy (Ψ) for each species. Estimates from our occupancy models are best interpreted as intensities of site use rather than true occupancy because our sampling units were smaller than the home range for species, so grid cells are not considered independent [27]. For all species, we modeled the detection process first using covariates expected to influence detection while holding occupancy constant. We then used the top detection model(s) to model species occupancy and incorporated variables among which occupancy may vary. We assessed overdispersion of the global model (*ĉ*) for each species and used QAICc model selection to account for overdispersion of the data. We determined the best model for each mammal species based on a model set of all combinations of covariates from a global model. We considered all models with ΔQAICc < 2 to be the best models describing species occupancy, and we chose the model with the lowest QAICc value to be the top performing model. We assessed the χ2 goodness-of-fit and produced QAICc model weights of each candidate model with ΔQAICc < 2.

We used the same detection variables in the global model for all species. We expected % savanna (SAV) to impact species’ detection, as open savanna habitat would improve visibility. We used the USGS land cover time series data from 2013 to extract the percentage of savanna habitat within our 10 x 10 km study grid cells [28]. We also expected variables related to the survey design to potentially impact the detection of all species: year (YR), camera type (CAM), and trap-nights (TN). Because we use different cameras, namely distinguishable by illumination specification, CAM was also a binary covariate representing white-flash and infrared models. TN, as a measure of sampling effort, was the total number of days that cameras were operational during survey. Lion (LIO), human (HUM), and livestock (LIVE) occupancy were included as covariates for all species and obtained from Mills *et al*. [26]. LIVE represent potential competitors and prey species for ungulates and hyena, respectively. Additionally, we incorporated competitor activity as trap success (detections per 100 trap-nights) of guild members of similar body size for ungulates and prey availability (ungulate detections per 100 trap-nights) for hyena. Ultimately, our interest lies in testing the effects of lion occupancy as a covariate explaining spatial patterns of competitor and prey species. We expect significant, negative beta coefficients for species, if lions are exerting top-down and intraguild pressures within the community. From the resultant occupancy estimates, we used a Mann-Whitney test to evaluate whether occupancy differed between low and high zones of lion activity. Again, we expect lower occupancy in high zones to indicate spatial avoidance, if the Critically Endangered West Africa lion population is still exerting regulatory pressures.

## 3. Results

Our camera survey yielded 96 independent lion detections based on aggregated images within a 30-minute quiet period. Lions exhibited spatial heterogeneity in their occupancy patterns across the WAP complex (figure 2a). Using the median occupancy (Ψ) estimate of 0.46 (range 0.25-1.0) as the threshold, we designated 88 and 116 grid cells of 10 km^2^ as low and high intensity of use by lions, respectively.

In comparison to lions, we obtained more than six times as many detections of hyenas (n=628), the most formidable competitor of lions [29]. Hyena occupancy varied from 0.001 – 1.0 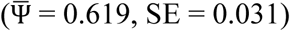 across the study area. Prey availability and lion occupancy both positively influenced hyena occupancy, as these two covariates repeatedly appeared in top models (table 1). We expected hyenas to exhibit a positive spatial response to prey availability because predators must follow food resources for their survival [e.g., 23, 30]. However, contrary to expectations, hyenas did not display a negative spatial response to lion presence; and instead, the pattern was opposite, with higher space use by hyenas in areas of heavy lion use. These results suggest lions are not exerting intraguild pressures on this competitor via spatial patterns, but radio-telemetry data would further strengthen this claim.

**Table 1.**
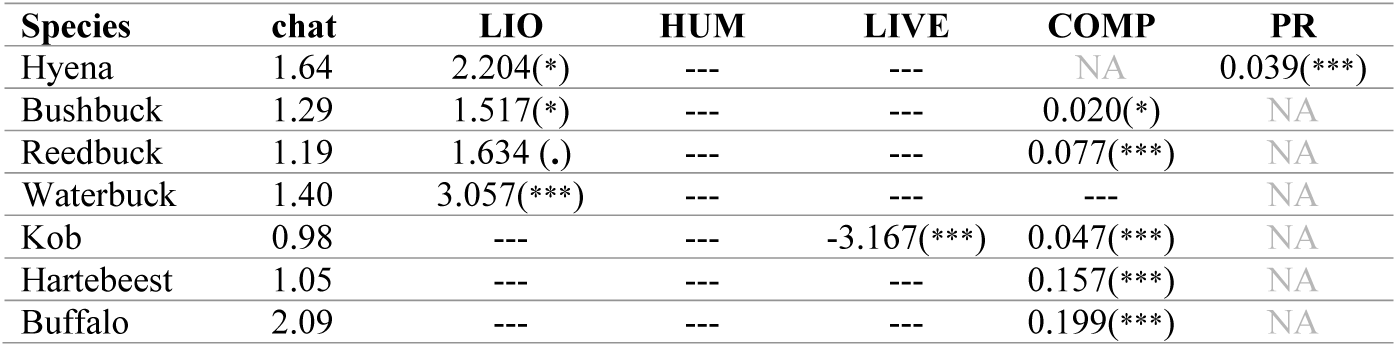
Beta coefficients from top occupancy models with overdispersion estimate (chat). Prey (PR) and competitors (COMP) only included for hyena and prey species, respectively. Only significant covariates are shown: (.) *p* < 0.1, (*) *p* < 0.05, (**) *p* < 0.01, (***) *p* < 0.001.

Prey species evaluated in our study included bushbuck (*Tragelaphus sylvaticus*, detections =2245), reedbuck (*Redunca redunca*, n=1270), kob (*Kobus kob*, n=847), buffalo (*Syncerus caffer*, n=698) and the less common hartebeest (*Alcelaphus buselaphus*, n=211) and waterbuck (*Kobus ellipsiprymnus*, n=161). If lions are not functionally extinct and are exerting top down pressures to regulate prey species, we expect these species to exhibit fear responses that manifest in avoidance behavior in the presence of lions. Many studies document spatial avoidance of risks across trophic levels including apex predators as sources of fear [e.g., 31, 32]. Within our occupancy-modeling framework, support for this hypothesis would result in negative spatial associations between the African lion and their ungulate prey. We observed significant associations between lion occupancy and that of three prey species but, contrary to expectation, these associations were positive (table 1, S1; reedbuck:β = 1.634, *p* = 0.089; bushbuck:β = 1.517, *p* = 0.040, and waterbuck (β = 3.057, *p* < 0.001).

Additionally, we evaluated whether significant differences in species occupancy for hyenas and ungulates existed across intensity use zones of lions. Again, contrary to expectations of lion avoidance, we found that for six of our seven focal species, occupancy actually increased in areas with high lion activity (figure 2b). Specifically, we observed significant increases for hyena (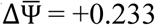, Mann-Whitney *p-value* < 0.001) and three prey species: waterbuck (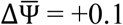, *p* = 0.007), reedbuck (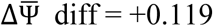, *p* = 0.007), and kob (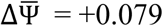, *p* = 0.018). The buffalo was the only species that exhibited a negative trend with higher occupancy in areas of low lion intensity (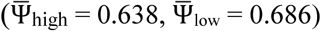), as expected; though, this difference was not significant.

## 4. Discussion

Our results raise concerns that lions have ceased to function as important regulators of ecological communities in a fragile West African ecosystem. We found no evidence that lions negatively influenced space use in competitor or common ungulate prey species within the WAP complex of West Africa. However, we recognize, as indicated with other covariates included in occupancy models, that the spatial distribution of these sympatric species cannot solely be attributed to lion space use. Similarly, M’soka *et al*. [33] found no effect of lions on hyena survival in a Zambian study area where lions have been nearly expirated. By contrast, in South Africa where lions persist at higher densities, Thaker *et al*. [32] reported low spatial overlap between lions and other carnivores, and all six ungulate species studied exhibited negative spatial responses to the presence of lions, even for species that were not primary prey for lions. Additionally, buffalo (*Syncerus caffer*) avoided “risky” waterholes and times as a defense against their primary lion predators in Zimbabwe where lions are not critically endangered as in comparison to our West African system [34]. Where lions become rare (e.g., critically endangered or endangered), other sympatric predators including hyenas and even humans may exert more pressure to structure and regulate mammal communities. We did not find evidence for humans ascending to apex predator status based on occupancy modeling results (Table S1). Lions compete with hyenas across their range with similar feeding patterns and studies have shown that prey species display comparable evasion strategies to hyenas as with lions [29]. Future work investigating whether hyenas ascend trophic levels where lions are locally extinct or have significantly dwindled would elucidate whether such a compensatory mechanism exists. Furthermore, we recommend increased financial and research investment focused on lion populations in West Africa to help restore this predator’s critical ecosystem function.

In the future, identifying other ecological indicators that signal the status of African lions would prove useful in monitoring and designing effective interventions in a timely manner. Here, we used spatial avoidance via occupancy models as the indicator. More nuanced aspects of the behavior of prey and competitor species could also be indicative of lion status garnered from experimental, physiological, and invasive studies. For example, prey might not alter spatial patterns, but instead increase group sizes and vigilance behavior as antipredator responses to African lions [35]. As such, these and other ecological indicators including the diversity and composition of parasite communities, fear responses on diel activity, and meat consumption of omnivores might signal functional extinction of an apex predator. We suspect that when researchers fail to detect effects of African lions on behavioral attributes and community structure, this may serve as a warning that lions are not functioning properly in the ecosystem and the population is at risk of extirpation.

Very few species couple natural and human ecosystems in the way that African lions do. Ecologically, lions alter nutrient pathways as well as modify behaviors in sympatric species [18, 19, 23]. Socially, lions contribute to economies through ecotourism and trophy hunting [36, 37] and compromise livelihoods for local people through depredation of domestic species that creates human-lion conflict [38, 39]. Given the occurrence of African lions has such broad implications for environment and society, research highlighting only the status and distribution of lions is therefore no longer sufficient. We underscore the necessity of scrutinizing their dynamic and perhaps dwindling functions within communities.

## Acknowledgements

We thank the AWE Lab for assistance with image identification and data management. We also appreciate feedback provided by R. Malhotra, N. H. Carter, A. King, and R. R. Dunn on earlier versions of the manuscript. We extend our sincerest appreciation to all park managers and the field team especially I. Gnoumou and Y.I. Abdel-Nasser that assisted with data collection and logistical support. We also thank administrators in Ministries of Environment in Burkina Faso (OFINAP & DGEF) and Niger (DGE/EF) especially B. Doamba and Y. Harissou for permitting and assisting with field management as well as private concessionaires in Burkina Faso for wildlife management efforts and access to properties in WAP. We also acknowledge University of Michigan (UM) African Studies Center - STEM initiative, UM Office of Research, the German Society of Mammalian Biology, and the Detroit Zoological Society for financial support.

## References

[1] Lindsey, P.A., Petracca, L.S., Funston, P.J., Bauer, H., Dickman, A., Everatt, K., Flyman, M., Henschel, P., Hinks, A.E., Kasiki, S., et al. 2017 The performance of African protected areas for lions and their prey. Biol. Conserv. 209, 137-149. (doi: http://dx.doi.org/10.1016/j.biocon.2017.01.011).

[2] Bauer, H., Chapron, G., Nowell, K., Henschel, P., Funston, P., Hunter, L.T.B., Macdonald, D.W. & Packer, C. 2015 Lion (Panthera leo) populations are declining rapidly across Africa, except in intensively managed areas. Proc. Natl. Acad. Sci 112, 14894-14899. (doi: 10.1073/pnas.1500664112).

[3] Henschel, P., Coad, L., Burton, C., Chataigner, B., Dunn, A., MacDonald, D., Saidu, Y. & Hunter, L.T.B. 2014 The Lion in West Africa Is Critically Endangered. Plos One 9, e83500. (doi: 10.1371/journal.pone.0083500).

[4] Hoag, C. & Svenning, J.C. 2017 African Environmental Change from the Pleistocene to the Anthropocene. Annual Review of Environment and Resources 42, 27-54. (doi: 10.1146/annurev-environ-102016-060653).

[5] Canning, D., Raja, S. & Yazbeck, A.S. 2015 Africa’s demographic transition: Dividend or disaster? (World Bank Publications.

[6] Thornton, P.K. 2010 Livestock production: recent trends, future prospects. Philosophical Transactions of the Royal Society B: Biological Sciences 365, 2853-2867. (doi: 10.1098/rstb.2010.0134).

[7] Carter, N.H., Bouley, P., Moore, S., Poulos, M., Bouyer, J. & Pimm, S.L. 2018 Climate change, disease range shifts, and the future of the Africa lion. Conserv. Biol. 32, 1207-1210. (doi: doi:10.1111/cobi.13102).

[8] Estes, J.A., Terborgh, J., Brashares, J.S., Power, M.E., Berger, J., Bond, W.J., Carpenter, S.R., Essington, T.E., Holt, R.D., Jackson, J.B.C., et al. 2011 Trophic Downgrading of Planet Earth. Science 333, 301-306. (doi: 10.1126/science.1205106).

[9] Ripple, W.J., Estes, J.A., Beschta, R.L., Wilmers, C.C., Ritchie, E.G., Hebblewhite, M., Berger, J., Elmhagen, B., Letnic, M., Nelson, M.P., et al. 2014 Status and Ecological Effects of the World’s Largest Carnivores. Science 343, 1241484. (doi: 10.1126/science.1241484).

[10] Heithaus, M.R., Frid, A., Wirsing, A.J. & Worm, B. 2008 Predicting ecological consequences of marine top predator declines. Trends Ecol. Evol. 23, 202-210. (doi: 10.1016/j.tree.2008.01.003).

[11] Packer, C., Holt, R.D., Hudson, P.J., Lafferty, K.D. & Dobson, A.P. 2003 Keeping the herds healthy and alert: implications of predator control for infectious disease. Ecol. Lett. 6, 797-802. (doi: 10.1046/j.1461-0248.2003.00500.x).

[12] Atkins, J.L., Long, R.A., Pansu, J., Daskin, J.H., Potter, A.B., Stalmans, M.E., Tarnita, C.E. & Pringle, R.M. 2019 Cascading impacts of large-carnivore extirpation in an African ecosystem Science 364, 173–177.

[13] Schmitz, O.J., Hawlena, D. & Trussell, G.C. 2010 Predator control of ecosystem nutrient dynamics. Ecol. Lett. 13, 1199-1209. (doi: 10.1111/j.1461-0248.2010.01511.x).

[14] Lima, S.L. 1998 Nonlethal effects in the ecology of predator-prey interactions. Bioscience 48, 25–34.

[15] Gaynor, K.M., Brown, J.S., Middleton, A.D., Power, M.E. & Brashares, J.S. 2019 Landscapes of Fear: Spatial Patterns of Risk Perception and Response. Trends Ecol. Evol. 34, 355-368. (doi: https://doi.org/10.1016/j.tree.2019.01.004).

[16] Moll, R.J., Killion, A.K., Montgomery, R.A., Tambling, C.J. & Hayward, M.W. 2016 Spatial patterns of African ungulate aggregation reveal complex but limited risk effects from reintroduced carnivores. Ecology 97, 1123-1134. (doi: 10.1890/15-0707.1).

[17] Owen-Smith, N. 2019 Ramifying effects of the risk of predation on African multi-predator, multiprey large-mammal assemblages and the conservation implications. Biol. Conserv. 232, 51-58. (doi: https://doi.org/10.1016/j.biocon.2019.01.027).

[18] Barnier, F., Valeix, M., Duncan, P., Chamaillé-Jammes, S., Barre, P., Loveridge, A.J., Macdonald, D.W. & Fritz, H. 2014 Diet quality in a wild grazer declines under the threat of an ambush predator. Proceedings of the Royal Society of London B: Biological Sciences 281. (doi: 10.1098/rspb.2014.0446).

[19] Le Roux, E., Kerley, G.I. & Cromsigt, J.P. 2018 Megaherbivores modify trophic cascades triggered by fear of predation in an African savanna ecosystem. Curr. Biol. 28, 2493–2499.

[20] Ford, A.T., Goheen, J.R., Otieno, T.O., Bidner, L., Isbell, L.A., Palmer, T.M., Ward, D., Woodroffe, R. & Pringle, R.M. 2014 Large carnivores make savanna tree communities less thorny. Science 346, 346-349. (doi: 10.1126/science.1252753).

[21] Schmitz, O.J., Wilmers, C.C., Leroux, S.J., Doughty, C.E., Atwood, T.B., Galetti, M., Davies, A.B. & Goetz, S.J. 2018 Animals and the zoogeochemistry of the carbon cycle. Science 362, eaar3213. (doi: 10.1126/science.aar3213).

[22] Dröge, E., Creel, S., Becker, M.S. & M’soka, J. 2017 Spatial and temporal avoidance of risk within a large carnivore guild. Ecology and Evolution 7, 189-199. (doi: 10.1002/ece3.2616).

[23] Vanak, A.T., Fortin, D., Thaker, M., Ogden, M., Owen, C., Greatwood, S. & Slotow, R. 2013 Moving to stay in place:behavioral mechanisms for coexistance of African large carnivores Ecology 94, 2619–2631.

[24] Hilborn, A., Pettorelli, N., Caro, T., Kelly, M.J., Laurenson, M.K. & Durant, S.M. 2018 Cheetahs modify their prey handling behavior depending on risks from top predators. Behav. Ecol. Sociobiol. 72. (doi: 10.1007/s00265-018-2481-y).

[25] Harris, N.C., Mills, K.L., Harissou, Y., Hema, E.M., Gnoumou, I.T., VanZoeren, J., Abdel-Nasser, Y.I. & Doamba, B. 2019 First camera survey in Burkina Faso and Niger reveals human pressures on mammal communities within the largest protected area complex in West Africa. Conservation Letters 12, e12667. (doi: 10.1111/conl.12667).

[26] Mills, K.L., Harissou, Y., Gnoumou, I.T., Abdel-Nasser, Y.I. Doamba & Harris, N.C. 2020 Comparable space use by lions between hunting concessions and national parks in West Africa. J. Appl. Ecol. 57, 975–984.

[27] MacKenzie, D.I., Bailey, L.L. & Nichols, J.D. 2004 Investigating species co-occurrence patterns when species are detected imperfectly. J. Anim. Ecol. 73, 546–555.

[28] Tappan, G.G., Cushing, W.M., Cotillon, S.E., Mathis, M.L., Hutchinson, J.A. & Dalsted, K.J. 2016 Data from: West Africa land use land cover time series. (ed. U.S.G.S.d. release.).

[29] Periquet, S., Fritz, H. & Revilla, E. 2015 The Lion King and the Hyaena Queen: large carnivore interactions and coexistence. Biological Reviews 90, 1197-1214. (doi: 10.1111/brv.12152).

[30] Burton, A.C., Sam, M.K., Balangtaa, C. & Brashares, J.S. 2012 Hierarchical Multi-Species Modeling of Carnivore Responses to Hunting, Habitat and Prey in a West African Protected Area. Plos One 7, e38007.

[31] Swanson, A., Caro, T., Davies-Mostert, H., Mills, M.G.L., Macdonald, D.W., Borner, M., Masenga, E. & Packer, C. 2014 Cheetahs and wild dogs show contrasting patterns of suppression by lions. J. Anim. Ecol. 83, 1418-1427. (doi: 10.1111/1365-2656.12231).

[32] Thaker, M., Vanak, A.T., Owen, C.R., Ogden, M.B., Niemann, S.M. & Slotow, R. 2011 Minimizing predation risk in a landscape of multiple predators: effects on the spatial distribution of African ungulates. Ecology 92, 398–407.

[33] M’Soka, J., Creel, S., Becker, M.S. & Droge, E. 2016 Spotted hyaena survival and density in a lion depleted ecosystem: The effects of prey availability, humans and competition between large carnivores in African savannahs. Biol. Conserv. 201, 348-355. (doi: http://dx.doi.org/10.1016/j.biocon.2016.07.011).

[34] Valeix, M., Fritz, H., Loveridge, A.J., Davidson, Z., Hunt, J.E., Murindagomo, F. & Macdonald, D.W. 2009 Does the risk of encountering lions influence African herbivore behaviour at waterholes? Behav. Ecol. Sociobiol. 63, 1483-1494. (doi: 10.1007/s00265-009-0760-3).

[35] Creel, S., Schuette, P. & Christianson, D. 2014 Effects of predation risk on group size, vigilance, and foraging behavior in an African ungulate community. Behav. Ecol. 25, 773-784. (doi: 10.1093/beheco/aru050).

[36] Lindsey, P.A., Balme, G.A., Booth, V.R. & Midlane, N. 2012 The Significance of African Lions for the Financial Viability of Trophy Hunting and the Maintenance of Wild Land. Plos One 7. (doi: 10.1371/journal.pone.0029332).

[37] Macdonald, D.W., Loveridge, A.J., Dickman, A., Johnson, P.J., Jacobsen, K.S. & Du Preez, B. 2017 Lions, trophy hunting and beyond: knowledge gaps and why they matter. Mamm. Rev. 47, 247-253. (doi: 10.1111/mam.12096).

[38] Ontiri, E.M., Odino, M., Kasanga, A., Kahumbu, P., Robinson, L.W., Currie, T. & Hodgson, D.J. 2019 Maasai pastoralists kill lions in retaliation for depredation of livestock by lions. People and Nature 1, 59-69. (doi: 10.1002/pan3.10).

[39] Hemson, G., Maclennan, S., Mills, G., Johnson, P. & Macdonald, D. 2009 Community, lions, livestock and money: a spatial and social analysis of attitudes to wildlife and the conservation value of tourism in a human–carnivore conflict in Botswana. Biol. Conserv. 142, 2718–2725.

