## Supplementary material for "Fear not: avoidance behavior missing in species sympatric with the critically endangered West African lion": Table S1

Supplementary Materials

### Do Critically Endangered West African lions still exert top-down pressures?

#### Supplemental Results

Table S1 Final candidate model set of all hyena and prey species occupancy models of ΔQAICc < 2 compared with the top performing model for each group. The bolded model indicates the top model used for comparison with lion presence. CAM = camera type, TN = trap-nights, YR = survey year, SAV = % savanna habitat, HUM = human occupancy, LIV = livestock occupancy, LN = lion occupancy, COMP = competitor species, PR = prey trap success per 100 trapnights.

| **Candidate Models** | **QAICc** | **ΔQAICc** | **QAICc weight** |
|---|---|---|---|
| ***HYENA (Competitor)*** | | | |
| *p* (CAM + TN + YR) *ψ* (LN + LIV + PR) | 795.92 | 0.16 | 0.24 |
| *p* (CAM + TN + YR) *ψ* (LIV + PR) | 796.69 | 0.94 | 0.16 |
| *p* (CAM + TN + YR) *ψ* (PR) | 797.01 | 1.25 | 0.14 |
| *p* (CAM + TN + YR) *ψ* (HUM + LN + LIV + PR) | 797.54 | 1.78 | 0.11 |
| *p* (CAM + TN + YR) *ψ* (HUM + LN + PR) | 797.62 | 1.86 | 0.10 |
| ***BUSHBUCK (Prey)*** | | | |
| ***p* (CAM + SAV) *ψ* (COMP + LN)** | **1245.45** | **0.00** | **0.27** |
| *p* (CAM + SAV + TN) *ψ* (COMP + LN) | 1246.46 | 1.02 | 0.16 |
| *p* (CAM + SAV) *ψ* (COMP + LN + LIV) | 1246.49 | 1.05 | 0.16 |
| *p* (CAM + SAV + YR) *ψ* (COMP + LIO) | 1246.78 | 1.33 | 0.14 |
| *p* (CAM + SAV) *ψ* (COMP) | 1246.81 | 1.37 | 0.14 |
| *p* (CAM + SAV) *ψ* (COMP + HUM + LIO) | 1246.84 | 1.39 | 0.13 |
| ***KOB (Prey)*** | | | |
| ***p* (CAM + SAV) *ψ* (COMP + LIV)** | **601.97** | **0.00** | **0.40** |
| *p* (CAM + SAV) *ψ* (HUM + COMP + LIV) | 602.81 | 0.84 | 0.26 |
| *p* (CAM + SAV + TN) *ψ* (COMP + LIV) | 603.71 | 1.74 | 0.17 |
| *p* (SAV) *ψ* (COMP + LIV) | 603.73 | 1.76 | 0.17 |
| ***REEDBUCK (Prey)*** | | | |
| ***p* (CAM + SAV) *ψ* (LN + COMP)** | **1264.55** | **0.00** | **0.12** |

| | | | |
|---|---|---|---|
| *p* (CAM + SAV + TN + YR) *ψ* (LN + COMP) | 1264.81 | 0.25 | 0.10 |
| *p* (CAM + SAV + TN) *ψ* (LN + COMP) | 1264.93 | 0.37 | 0.10 |
| *p* (CAM + SAV) *ψ* (COMP) | 1265.23 | 0.67 | 0.08 |
| *p* (CAM + SAV) *ψ* (LN + LIV + COMP) | 1265.30 | 0.75 | 0.08 |
| *p* (CAM + SAV + TN + YR) *ψ* (COMP) | 1265.51 | 0.96 | 0.07 |
| *p* (CAM + SAV + TN) *ψ* (COMP) | 1265.55 | 0.99 | 0.07 |
| *p* (CAM + SAV + TN) *ψ* (LN + LIV + COMP) | 1265.72 | 1.17 | 0.06 |
| *p* (CAM + SAV + YR) *ψ* (LN + COMP) | 1265.75 | 1.19 | 0.06 |
| *p* (CAM + SAV) *ψ* (LIV + COMP) | 1265.87 | 1.32 | 0.06 |
| *p* (CAM + SAV + TN + YR) *ψ* (LN + LIV + COMP) | 1265.88 | 1.33 | 0.06 |
| *p* (CAM + SAV + TN) *ψ* (LIV+ COMP) | 1266.24 | 1.69 | 0.05 |
| *p* (CAM + SAV + YR) *ψ* (COMP) | 1266.48 | 1.91 | 0.04 |
| *p* (CAM + SAV + TN + YR) *ψ* (LIV + COMP) | 1266.48 | 1.93 | 0.04 |
| ***WATERBUCK (Prey)*** | | | |
| ***p* (TN) *ψ* (LN)** | **299.54** | **0.00** | **0.51** |
| *p* (TN) *ψ* (LN + LIV) | 300.76 | 1.22 | 0.28 |
| *p* (SAV + TN) *ψ* (LN) | 301.33 | 1.80 | 0.21 |
| ***HARTEBEEST (Prey)*** | | | |
| **p (CAM + SAV + TN) *ψ* (COMP)** | **814.72** | **0.00** | **0.28** |
| p ( SAV + TN) *ψ* (COMP) | 814.96 | 0.25 | 0.25 |
| p (CAM + SAV + TN) *ψ* (COMP + HUM) | 815.02 | 0.31 | 0.24 |
| p ( SAV + TN) *ψ* (COMP + HUM) | 815.05 | 0.34 | 0.24 |
| ***BUFFALO (Prey)*** | | | |
| **p (YR) *ψ* (COMP)** | **660.67** | **0.00** | **0.20** |
| p (CAM) *ψ* (COMP) | 660.77 | 0.09 | 0.19 |
| p (.) *ψ* (COMP) | 661.69 | 1.02 | 0.12 |
| p (SAV + YR) *ψ* (COMP) | 661.94 | 1.27 | 0.10 |
| p (CAM + YR) *ψ* (COMP) | 662.23 | 1.56 | 0.09 |
| p (CAM+ TN) *ψ* (COMP) | 662.26 | 1.59 | 0.09 |
| p (TN) *ψ* (COMP) | 662.56 | 1.89 | 0.08 |
| p (TN + YR) *ψ* (COMP) | 662.65 | 1.98 | 0.07 |

| | | | |
|---|---|---|---|
| p (CAM + SAV) $\psi$ (COMP) | 662.66 | 1.99 | 0.07 |
